# Prenatal Inflammation as a Determinant of Long-Term Cardiovascular Risk: Evidence from a Murine Model

**DOI:** 10.64898/2026.09.15.750739

**Authors:** K. Kersh, M. Patel, J. Roble, A.C. Aminkeng, M. Rath, E. Bytautiene Prewit

## Abstract

**Background:** Intrauterine inflammation is associated with increased lifetime cardiovascular disease (CVD) risk in offspring, yet long-term molecular mechanisms remain underexplored. We investigated the lasting cardiometabolic, hemodynamic, and vascular effects of prenatal inflammatory exposure in adult offspring using a mouse model.

**Methods:** On day 15 of gestation, pregnant CD-1 mice were randomized to receive intraperitoneal (IP) injections of either lipopolysaccharide (LPS group) or saline (SAL group). Blood samples were collected before and after injection to measure IL-6 concentrations. Blood pressure (BP) in offspring was measured at 6-8 months of age using the tail-cuff method, and tissues were collected. Protein was extracted from the brain cortex to measure cytokines utilizing the xMAP Luminex assay. The C-reactive protein (CRP) was measured in serum. Gene expression of IL-6, NOS3, NFkB, AT-1, and Trp53 was determined by RT-qPCR in the aorta. The experimental unit was a mother. Results were analyzed using the appropriate statistical methods.

**Results:** Maternal LPS administration induced significant acute systemic inflammation (P<0.05 for IL-6). At 6-8 months after delivery, maternal circulating CRP was significantly higher in LPS mice as compared to SAL (P=0.02). In offspring at 6-8 months of age, systolic BP was significantly elevated in LPS groups compared with SAL animals (P=0.04). LPS-exposed offspring exhibited higher total body and visceral adipose tissue weights. Aortic expression of the angiotensin II type 1 receptor (AT1) and NFkB were significantly upregulated (P=0.04 and P=0.03, respectively). Cortices of LPS offspring displayed a pro-inflammatory profile with increased total Inflammatory score, elevated pro-inflammatory IL-1β, and reduced anti-inflammatory IL-4 and IL-10. Furthermore, protective Trp53 gene expression was significantly downregulated in both the aorta (P=0.01) and heart (P=0.04).

**Conclusions:** A single mid-gestation inflammatory insult programs long-term cardio-inflammatory perturbations in offspring and results in long-term systemic inflammation in mothers. Prenatal LPS exposure induces persistent visceral obesity, hypertension, vascular RAS upregulation, central neuroinflammation, and cardiac/vascular Trp53 suppression in adult offspring, offering a mechanistic link between prenatal inflammation and offspring CVD risk.

## INTRODUCTION

Cardiovascular disease (CVD) is the leading cause of death in the US and worldwide (National Center for Health Statistics Mortality Data on CDC WONDER). Adverse pregnancy outcomes have been linked to an increased risk for CVD in both mothers and offspring. This association arises from vascular abnormalities and placental dysfunction, which can result in inflammation, hypertension, and other complications^1^.

While controlled inflammation is essential for trophoblastic invasion and remodeling of maternal blood vessels to ensure sufficient blood flow and nutrients to the fetus, exaggerated prenatal inflammation has become increasingly recognized as a critical factor affecting maternal health, fetal outcome, and development of diseases^2^. Excessive inflammation, as characterized by overactivation of inflammatory pathways, can compromise placental function and lead to adverse pregnancy outcomes including preterm birth, preeclampsia, low birth weight, and gestational diabetes^2^. Intrauterine inflammation is histologically present in 20% of the population, but this figure rises to 85% in cases of very preterm births^3^. Higher levels of pro-inflammatory cytokines correlate with the onset of preterm labor, while chronic maternal inflammation can interfere with placental function and hinder fetal growth^4^. Moreover, inflammatory mediators can impact the developing fetal brain, potentially leading to long-term neurodevelopmental, cognitive, and behavioral changes^5^.

Preterm births may predispose mothers and offspring to a range of adverse health conditions including cardiovascular disease, metabolic disorders, and psychological sequelae. In a meta-analysis of four studies, preterm birth was significantly linked to CVD mortality in women^6^. Women were at higher risk of developing chronic hypertension, type 2 diabetes, and hypercholesterolemia in the years following pregnancy^7^. Prenatal inflammation has also been associated with negative cardiovascular outcomes in offspring. A recent systematic review and meta-analysis of 28 studies found a significant link between preterm birth and hypertension in adulthood^8^. This intergenerational transmission of risk highlights the importance of understanding the mechanisms through which prenatal inflammation and preterm birth elevated cardiovascular risks of both mothers and offspring. Animal models have been instrumental in characterizing the inflammatory mechanisms driving preterm birth and its downstream consequences. A systematic review of 24 mouse model studies of infection and inflammation-induced preterm birth described that LPS exposure reliably recapitulates the inflammatory endotype of human preterm birth, validating its use as a preclinical platform for studying prenatal inflammatory pathophysiology^9^. However, a majority of these studies have focused on acute gestational outcomes such as fetal survival, timing of delivery, and immediate inflammatory markers, leaving the long-term cardiovascular consequences in surviving offspring largely uncharacterized. This gap represents a critical limitation in the field, as the mechanisms by which prenatal inflammatory exposure programs lasting cardiovascular dysfunction in offspring remain poorly understood.

The objective of our study was to developed a mouse model to study the long-term cardiovascular outcomes in offspring exposed to exaggerated prenatal inflammation.

## MATERIALS AND METHODS

### Ethical Approval and Animal Housing

All experimental protocols were approved by the Institutional Animal Care and Use Committee (IACUC) at the University of Texas San Antonio. Male and mature cycling female CD-1 mice were purchased from Charles River Laboratories (Wilmington, MA). Animals were housed in temperature- and humidity-controlled quarters under a 12:12-hour light/dark cycle. Pregnant females were housed individually to minimize stress.

### Experimental Design & Prenatal Exposure

Mice were bred by caging one male with two to three female mice overnight. Day one of pregnancy was recorded as the day a vaginal plug was present. On day 15 of gestation, pregnant CD-1 mice were randomized to receive intraperitoneal (IP) injections of either lipopolysaccharide (LPS, 50 μg in 1 mL of saline) or saline (SAL, 1 mL). Mice received a single injection of either lipopolysaccharide (LPS) or saline (control). Two hours later, all animals received a second saline injection to standardize handling and injection procedures across groups and to control for potential effects associated with repeated handling and injection-related stress. Blood samples in pregnant females were collected before and 6 hours post-injection and allowed to deliver. Delivery occurred on day 20-21 of pregnancy. Weights, blood pressure, molecular biology, and blood assays were performed on offspring at 6-8 months of age and on mothers at 10-12 months of age, corresponding to 6-8 months postpartum.

### Morphometric and Hemodynamic Assessments

At 6–8 months of age, offspring were evaluated for morphometric and hemodynamic changes. Total body weight (TBW) and visceral adipose tissue (VAT) weight were measured. Blood pressure was measured in vivo using the CODA non-invasive tail-cuff plethysmography system (Kent Scientific, Torrington, CT). CODA has been validated with telemetry with 99% correlation, as described by us previously^10^. Mice were acclimated to holders, and blood pressure was recorded over 20 continuous inflation/deflation cycles to obtain an averaged value for each animal.

### Biochemical Assays

Blood samples in pregnant females were collected before and hourly post-injection to measure circulating IL-6 concentrations. Protein was extracted from the brain cortex of the offspring to measure cytokine concentrations (IL-1 β, IL-4, IL-6, IL-10, TNFα) utilizing Mouse High Sensitivity T Cell Magnetic Bead Panel (cat. # MHSTCMAG-70K, Millipore, Billerica, MA, USA) according to the manufacturer’s instructions. The C-reactive protein (CRP) was measured in plasma (cat. # RAB1121, Sigma-Aldrich, St. Louis, MO, USA). Assays were performed in duplicate for each sample.

### RNA extraction and cDNA synthesis

Frozen aorta and heart tissue (50 mg) was homogenized in 500 μl TRIzol (Invitrogen) using a homogenizer (Next Advance Bullet Blender Storm ProBT24M). An RNeasy Mini Kit method (QIAGEN, Cat. No. 74106) was used to extract total RNA. The RNA concentration was measured using a Nanodrop™ Spectrophotometer (ND-1000). Total RNA for each sample was diluted to 500 ng/μl. Half of the total reaction volume (10 μl) was used for cDNA synthesis, and the rest was stored at −80°C. DNase-digested RNA (10 μl) was reverse-transcribed to cDNA using the Applied Bio Hi Capacity cDNA kit (Cat. No. 4368814). The following cycling conditions were used: 25° C for 10 min, 37° C for 120 min, 85° C for 5 min. The cDNA concentration for qPCR was calculated to be ~50ng/μl, under the assumption that 1–5% of total RNA is mRNA.

### Quantitative PCR

Samples of cDNA (~ 50 ng) were quantified by quantitative PCR using SYBR-Green detection methods in a Biorad CFX thermal cycler (BioRad, Hercules, CA). The primer sequences used were: NF-kB forward (5’-gctgccaaagaaggacacgaca-3’), NF-kB reverse (5’-ggcaggctattgctcatcacag-3’), NOS3 forward (5’-cgcaagaggaaggagtctagca-3’), and NOS3 reverse (5’-tcgagcaaaggcacagaagtgg-3’), AT-1 forward (5’-gccattgtccacccgatgaagt-3’), AT-1 reverse (5’-acacatttcggtggatgacggc-3’), IL-1β forward (5’-gcaactgttcctgaactcaact-3’), IL-1β reverse (5’-atcttttggggtccgtcaact-3’), IL-6 forward (5’-gcagcatcaccttcgcttaga-3’), IL-6 reverse (5’-cagatattggcatgggagcaag-3’), β-actin forward (5’-cggttccgatgccctgaggctctt-3’), β-actin reverse (5’-cgtcacacttcatgatggaattga-3’). Cycling conditions were 94 °C for 5 min, followed by 40 cycles of denaturation at 94 °C for 30 seconds and annealing at 60 °C for 30 seconds. Experimental plates were set up to include triplicates on the same run.

### Relative expression calculations

Raw Cq values from 3 technical replicates were used to calculate the mean Cq for the reference gene (β-actin) and the mean Cq for the genes of interest, NF-κB, NOS3, AT-1, IL-1β, and IL-6, for each sample. ΔCq were calculated as mean Cq(NF-KB) - mean Cq(β-actin), mean Cq(NOS3) - mean Cq(β-actin), Cq(AT-1) - mean Cq(β-actin), Cq(IL-1β) - mean Cq(β-actin),Cq(IL-6) - mean Cq(β-actin) or each sample. Relative expression was calculated as 2^- (ΔΔCt).

### Statistical Analysis

The experimental unit in this study was a mother (n=3 in LPS, n=3 in SAL group). A composite Inflammatory Index for the net inflammatory balance determination was calculated for each experimental unit as z(IL-1β) + z(IL-6) + z(TNFα) – z(IL-4) – z(IL-10). Statistical analysis was performed using GraphPad Prism version 11.1.0 for Mac OS X, GraphPad Software, Boston, Massachusetts, USA, www.graphpad.com. Data were tested for outliers, and then, depending on normality and differences between variances, Student’s t-test, Mann-Whitney test, or ordinary one-way ANOVA (with appropriate post-hoc tests) were used. A P value <0.05 was considered statistically significant. Results are presented as violin plots, unless stated otherwise. For violin plot we used medium smoothing, while the lines represent median and quartiles.

## RESULTS

On day 15 of gestation, circulating IL-6 levels in mothers were significantly higher in the LPS group at 6 hours after LPS injection when compared to the IL-6 concentration in SAL and LPS groups before the injection and in SAL group at 6 hours after injection (Fig. 1, P=0.01 for all comparisons).

**Fig 1.**
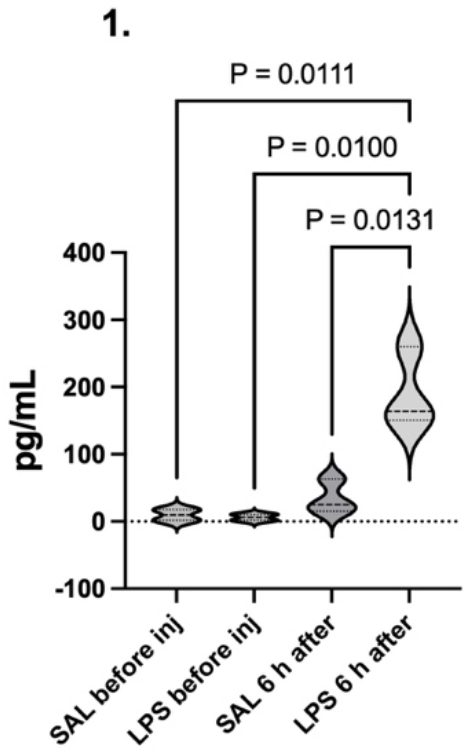
Concentration of IL-6 in maternal serum before the SAL (SAL before inj) or LPS (LPS before inj) injection and 6 hours after SAL (SAL 6 h after) or LPS (LPS 6 h after) injection.

Total body and visceral adipose tissue weights and adiposity (percent of visceral adipose tissue from total body weight) did not differ between the offspring born to dams after LPS administration and offspring of dams given SAL (data not shown).

The systolic BP in offspring born to LPS mothers was significantly higher compared to offspring born to SAL mothers (Fig. 2, P=0.04). Diastolic BP and mean BP were also higher in the LPS group, but were not statistically different (P=0.10 and P=0.07, respectively; data not shown). Protein levels in the offspring’s brain cortices were measured as mean fluorescence intensity (MFI) using multiplex antibody-coated bead xMAP Luminex. Of the proteins measured, the MFI of IL-1β was increased (P=0.16, Fig. 3A), while both IL-4 (P= 0.25, Fig. 3B) and IL-10 (P= 0.24, Fig. 3C) were lower in offspring exposed to LPS than in offspring from the SAL group, though these differences were not statistically significant. The inflammatory index was higher in the LPS group but did not reach statistical significance (P=0.07, Fig. 3D).

**Fig 2.**
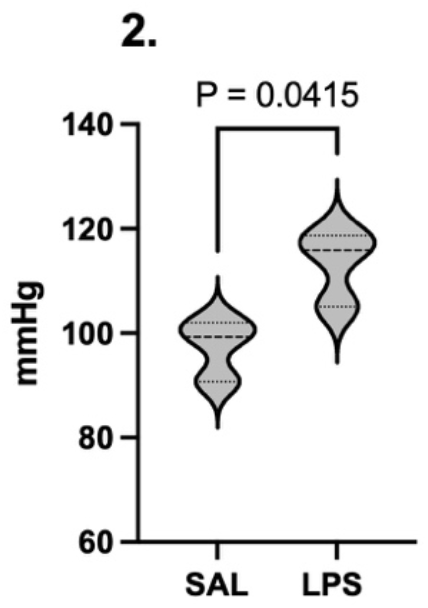
Systolic blood pressure of offspring born to mothers with prenatal administration of either SAL or LPS at 6-8 months of age.

**Fig 3.**
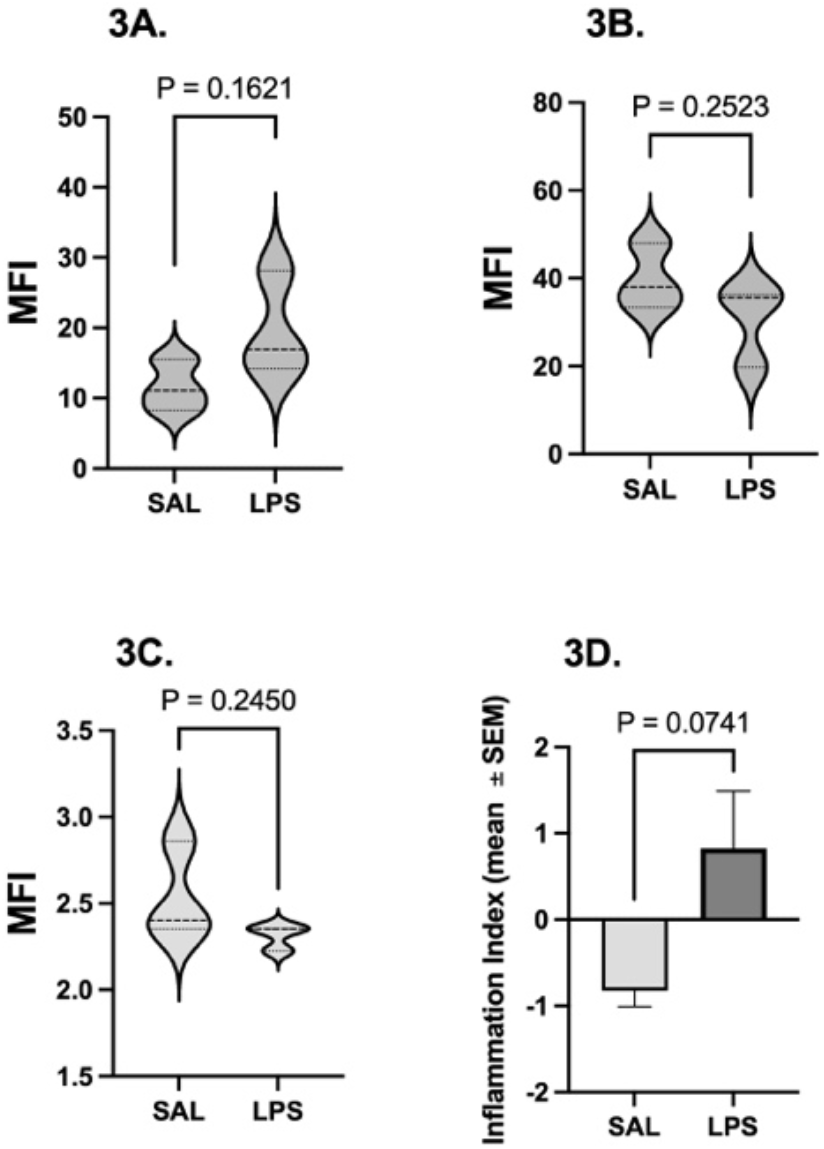
Mean fluorescence intensity (MFI) indicating protein levels of IL-1β (A), IL-4 (B), and IL-10 (C) and Inflammatory Index (D) in the cortex of offspring born to mothers with prenatal administration of either SAL or LPS at 6-8 months of age. Inflammtory Index for both groups is presented as mean ± SEM.

The AT1 receptor and NF-κB gene expressions in the aorta of offspring born to mothers with prenatal administration of LPS were significantly higher than in the SAL group at 6-8 months of age (P=0.04, Fig. 4A and P=0.03, Fig. 4B, respectively). IL-6 expression increased, and NOS3 expression decreased in the LPS group, but these changes did not reach statistical significance (data not shown).

**Fig 4:**
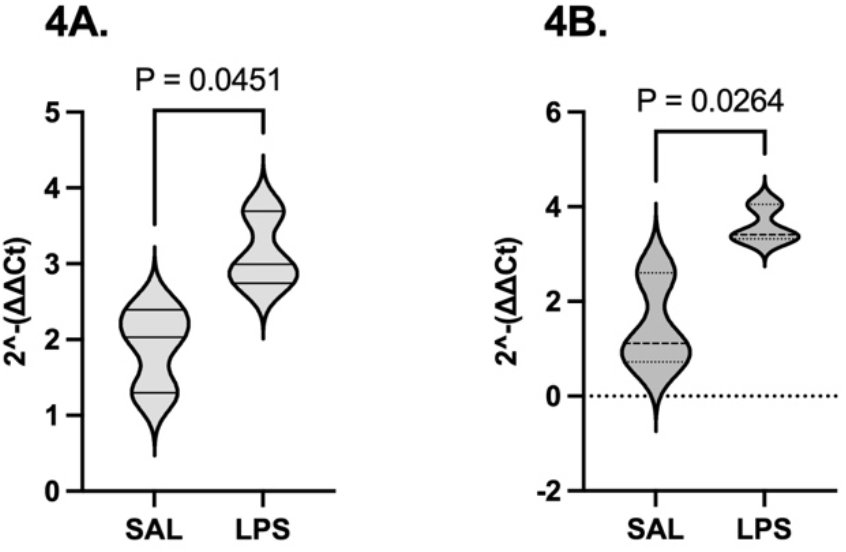
AT1 receptor (A) and NF-κB (B) expressions in the aorta of offspring born to mothers with prenatal administration of either SAL or LPS at 6-8 months of age.

TP53 expression in the aorta and heart of the offspring was significantly lower in the LPS group than in the SAL group (P=0.01, Fig. 5A; P=0.04, Fig. 5B), indicating impaired protection against inflammation in the LPS group.

**Fig 5.**
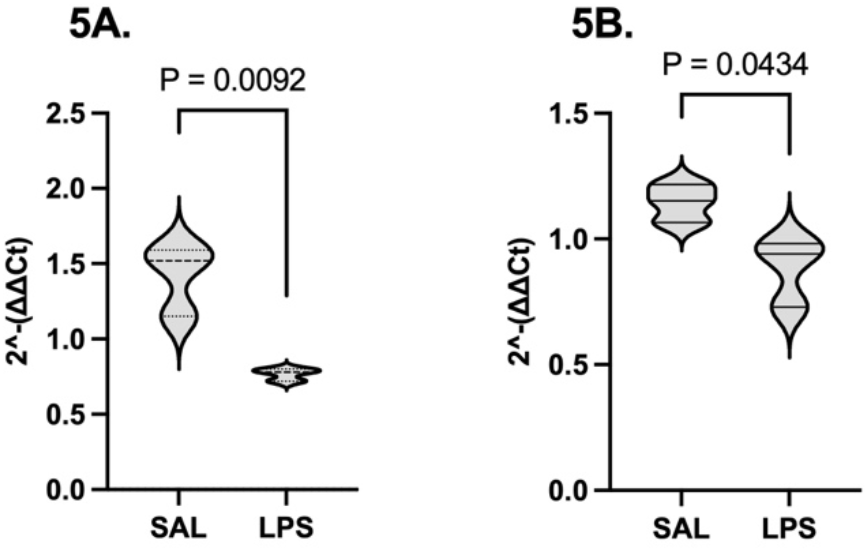
mRNA expression of Trp53 of offspring in the aorta (A) and heart (B) at 6-8 months of age.

We also measured CRP levels in mothers at 6-8 months post delivery. CRP was significanly higher in mothers that received LPS injection as compared to SAL group (P=0.02, Fig. 6).

**Fig 6.**
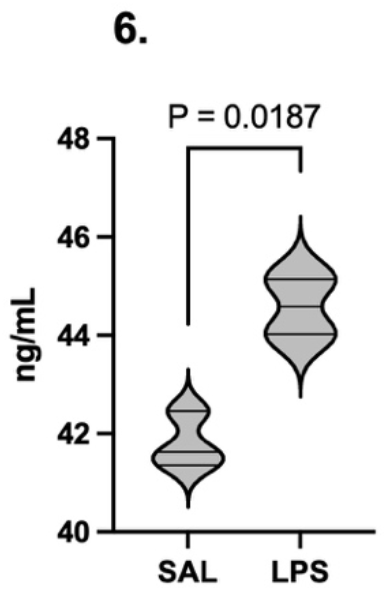
Circulating CRP concentration in mothers at 6-8 months after delivery.

## DISCUSSION

Our study establishes that a single, transient mid-gestational inflammatory insult can exert multi-systemic cardiorespiratory and metabolic programming effects that persist into mature adulthood in both offspring and mothers. In a murine model of inflammation-driven preterm birth, prenatal LPS exposure triggered acute systemic maternal-fetal inflammation and led to adult-onset hypertension, RAS hyperactivity, suppression of protective baseline Trp53 transcriptional activity, and sustained cortical neuroinflammation at 6–8 months of age. These findings suggest that prenatal inflammation alters developmental programming windows, inducing chronic vascular, metabolic, and neural dysfunction that spans across generations.

Administration of LPS on gestational day 15 induced a robust inflammatory surge, evidenced by marked systemic elevation of IL-6 when compared to saline controls. This cytokine response models human intra-amniotic infection, chorioamnionitis, and fetal inflammatory response syndrome (FIRS), which are major etiologic drivers of spontaneous preterm delivery^11-13^. Toll-like receptor 4 (TLR4) activation at the maternal-fetal interface initiates a signal transduction cascade that elevates pro-inflammatory cytokines in amniotic fluid and fetal circulation. Early inflammatory exposure disrupts critical windows of cellular differentiation and tissue organogenesis, laying the molecular groundwork for adult-onset chronic disease.

At 6–8 months of age, offspring born following prenatal LPS exposure exhibited greater total body weight and visceral adipose tissue mass compared to saline-exposed controls. This extends earlier work showing that prenatal LPS exposure increases offspring body weight and visceral fat mass within the first 8–12 weeks of life, suggesting that this impact on the metabolic syndrome is not transient but persists well into later life^14^. This pattern parallels observations in preterm-born humans, who exhibit a distinct cardiometabolic phenotype marked by increased adiposity, insulin resistance, and atypical growth trajectories that becomes evident beyond the neonatal perio^15,16^. Notably, this adiposity in preterm-born humans is preferentially visceral rather than subcutaneous, a distribution pattern consistent with what we observed in our LPS-exposed offspring^15^.

The most clinically significant finding in offspring was the presence of significantly elevated systolic blood pressure at 6–8 months of age in LPS-exposed animals. Meta-analyses of human cohort studies report that individuals born preterm carry an increased risk of hypertension and demonstrate approximately 2–5 mmHg higher systolic blood pressure compared to term-born peers^17^.

To elucidate the molecular basis of this hypertensive phenotype, we evaluated aortic gene expression and identified a significant upregulation of the AT1 receptor. Our data demonstrated that gestational inflammatory stress establishes long-term nuclear factor kappa B (NF-κB) dyshomeostasis in conduit arteries, which directly activates AT1 receptor expression^18^. Downstream activation of the vascular angiotensin II-AT1R axis stimulates membrane-bound NADPH oxidase complexes (NOX1 and NOX4), generating excessive intracellular reactive oxygen species (ROS). Excess superoxide radicals rapidly scavenge bioavailable nitric oxide (NO) to form peroxynitrite (ONOO−), inducing uncoupling of endothelial nitric oxide synthase (eNOS) and profound endothelial dysfunction.

Concurrently, systemic oxidative stress driven by prenatal inflammation suppresses renal dopamine D1 receptor (D1R) expression and phosphorylation, impairing renal tubular sodium excretion and diuresis^19^. In resistance vessels, augmented AT1R signaling triggers mitogen-activated protein kinase (MAPK) and extracellular signal-regulated kinase (ERK1/2) pathways, driving vascular smooth muscle cell (VSMC) hypertrophy, collagen deposition, and permanent arterial stiffening. These intracellular pathways lead to vascular remodeling, endothelial dysfunction, cardiovascular diseases, atherosclerosis, and end-organ damage^20^. LPS offspring at 6-8 months of age also showed increased Inflammatory Index, elevated IL-1β and reduced IL-4 and IL-10 in the cortex, a net pro-inflammatory phenotype. These findings suggest that prenatal insults affect multiple organ systems simultaneously, consistent with literature showing cytokines’ role in neurodevelopment in preterm births^21^. However, reduced IL-10 is particularly notable given its role in dampening vascular inflammation.

A particularly novel finding of this study is the significant downregulation of Trp53 in both the aorta and heart of LPS-exposed adult offspring. p53 is essential for embryonic cardiac development and is necessary to maintain normal heart architecture and physiological function^22^. Our study’s findings may reflect a loss of protective basal tone during a critical programming window, contributing to worse cardiovascular outcomes in offspring. Mothers who experienced LPS-induced inflammation demonstrated increased higher systemic inflammation at 6–8 months postpartum as evidenced by higher CRP levels in these dams. Systemic inflammation is a risk facot for many chronic dieseases. Our finding is consistent with a growing body of evidence of human data linking pregnancy complications (preeclampsia, preterm birth, pregnancy loss) to an elevated lifetime maternal risk of hypertension and other cardiovascular diseases^6,23–25^.

The key strengths of our study is that our results fill a gap in the literature characterizing adult-stage cardiovascular outcomes—evaluating metabolic, hemodynamic, neuroimmune, and vascular molecular markers at 6–8 months of age rather than focusing solely on acute gestational effects. This long-term framework connects mid-gestational inflammation to adult-onset disease. However, limitations must be noted. A single LPS injection models an acute endotoxin insult, which may not fully reflect the chronic or polymicrobial infectious phenotypes seen in human chorioamnionitis. Additionally, while we identified transcriptional changes like aortic AT1 receptor upregulation and Trp53 suppression, functional protein-level validation and targeted interventional studies are needed.

## CONCLUSION

In summary, our findings support a model of developmental programming wherein acute mid-gestational maternal inflammation initiates a cascade of persistent physiological alterations. These early inflammatory events lead to sustained RAS upregulation, loss of Trp53-mediated vascular protection, and neuroimmune dysregulation, ultimately converging on a phenotype of adult hypertension and visceral adiposity. This pathway provides a unifying framework linking prenatal inflammatory exposure to the elevated lifetime cardiovascular risk observed in clinical populations.

## Notes

Financial Disclosure: None of the authors have a conflict of interest

### Competing Interest Statement

The authors have declared no competing interest.

